# Herd protection against *Plasmodium falciparum* infections conferred by mass antimalarial drug administrations and the implications for malaria elimination

**DOI:** 10.1101/393843

**Authors:** Daniel M. Parker, Sai Thein Than Tun, Lisa J. White, Ladda Kajeechiwa, May Myo Thwin, Jordi Landier, Victor Chaumeau, Vincent Corbel, Arjen M. Dondorp, Lorenz von Seidlein, Nicholas J. White, Richard J. Maude, François H. Nosten

## Abstract

The global malaria burden has decreased over the last decade and many nations are attempting elimination. Asymptomatic infections aren’t normally diagnosed or treated, posing a major hurdle for elimination efforts. One solution to this problem is mass drug administration (MDA), which is dependent on adequate population participation to disrupt transmission. There is little empirical evidence regarding the necessary threshold level of participation. Here we present a detailed spatiotemporal analysis of malaria episodes and asymptomatic infections in four villages undergoing MDA in Myanmar. Individuals from neighborhoods with high MDA adherence had 90% decreased odds of having a malaria episode post-MDA, regardless of individual participation, suggesting a strong herd effect. High mosquito biting rates, living in a house with someone else with malaria, or having an asymptomatic malaria infection were also predictors of clinical episodes. Spatial clustering of non-adherence to MDA, even in villages with high overall participation, can frustrate elimination efforts.

## INTRODUCTION

Malaria in the Greater Mekong Subregion (GMS) is heterogeneous in both space and time (Cui, Yan, Sattabongkot, Cao, et al., 2012). Most symptomatic malaria is seasonal and the disease clusters along international borders (Cui, Yan, Sattabongkot, Cao, et al., 2012; Cui, Yan, Sattabongkot, Chen, et al., 2012; Parker, Carrara, Pukrittayakamee, Mcgready, & Nosten, 2015). All of the nations of the GMS have committed to eliminating malaria by 2030 (WHO, 2016) and cases of *Plasmodium falciparum* malaria have been decreasing over the last several decades. However, parasites are developing resistance to the first line of antimalarials (Phyo et al., 2016), presenting a public health emergency in the current absence of appropriate replacement therapies. One proposed solution to this problem is the elimination of *P. falciparum* parasites while prevalence is low and while combination therapies remain effective in clearing low density infections (von Seidlein & Dondorp, 2015).

The foundation of malaria elimination programs is the provision of access to early diagnosis and treatment (EDT) to all affected communities (Jordi Landier et al., 2016). Throughout the GMS this has been facilitated through programs which employ village or community health workers (Carrara et al., 2006); a system that has existed in parts of the GMS for over 50 years (especially in Thailand, Vietnam and China) (Kauffman & Myers, 1997; Lehmann & Sanders, 2007; Zhang & Unschuld, 2008). Infected and symptomatic individuals who quickly seek and receive early diagnosis and treatment have reduced infectious periods, leading to a reduction in transmission.

Recent work from several different parts of the GMS found several populations with high asymptomatic malaria prevalence (Imwong et al., 2015). Asymptomatic individuals are not treated and serve as a reservoir for persistence of malaria in the region even in the presence of strong healthcare systems. Therefore, another key tool in the elimination portfolio is targeted mass drug administration (MDA). MDA includes the administration of antimalarials to all individuals in a targeted population, in this case in communities that have high malaria prevalence (von Seidlein & Dondorp, 2015). There is likely to be a critical threshold for MDA coverage, below which the reduction of the parasite reservoir is not sufficient to halt ongoing transmission. However, there is little empirical evidence of such coverage effects.

Here we describe geographic and epidemiologic patterns of clinical and subclinical *P. falciparum* and *P. vivax* infections in villages receiving targeted MDA. We investigate associations between heterogeneous adherence to MDA, mosquito vector biting rates, asymptomatic infections, and clinical malaria episodes. To our knowledge this is the first subvillage level spatiotemporal analysis of malaria incidence and prevalence from an any MDA study.

## Methods

### Study location and design

The study site consisted of four villages (KNH, TPN, HKT, and TOT) along the Myanmar-Thailand border, in Kayin (Karen) State, Myanmar (Jordi Landier et al., 2017). The villages were selected based on malaria prevalence estimates from a preliminary survey in the area (Imwong et al., 2015) and were part of a MDA pilot study (Jordi Landier et al., 2017). The northernmost village is approximately 105 km from the southernmost and the two closest villages, KNH and TPN are within 10km of each other (**Figure 1**). The study was conducted from May 2013 through June 2015.

**Figure 1:**
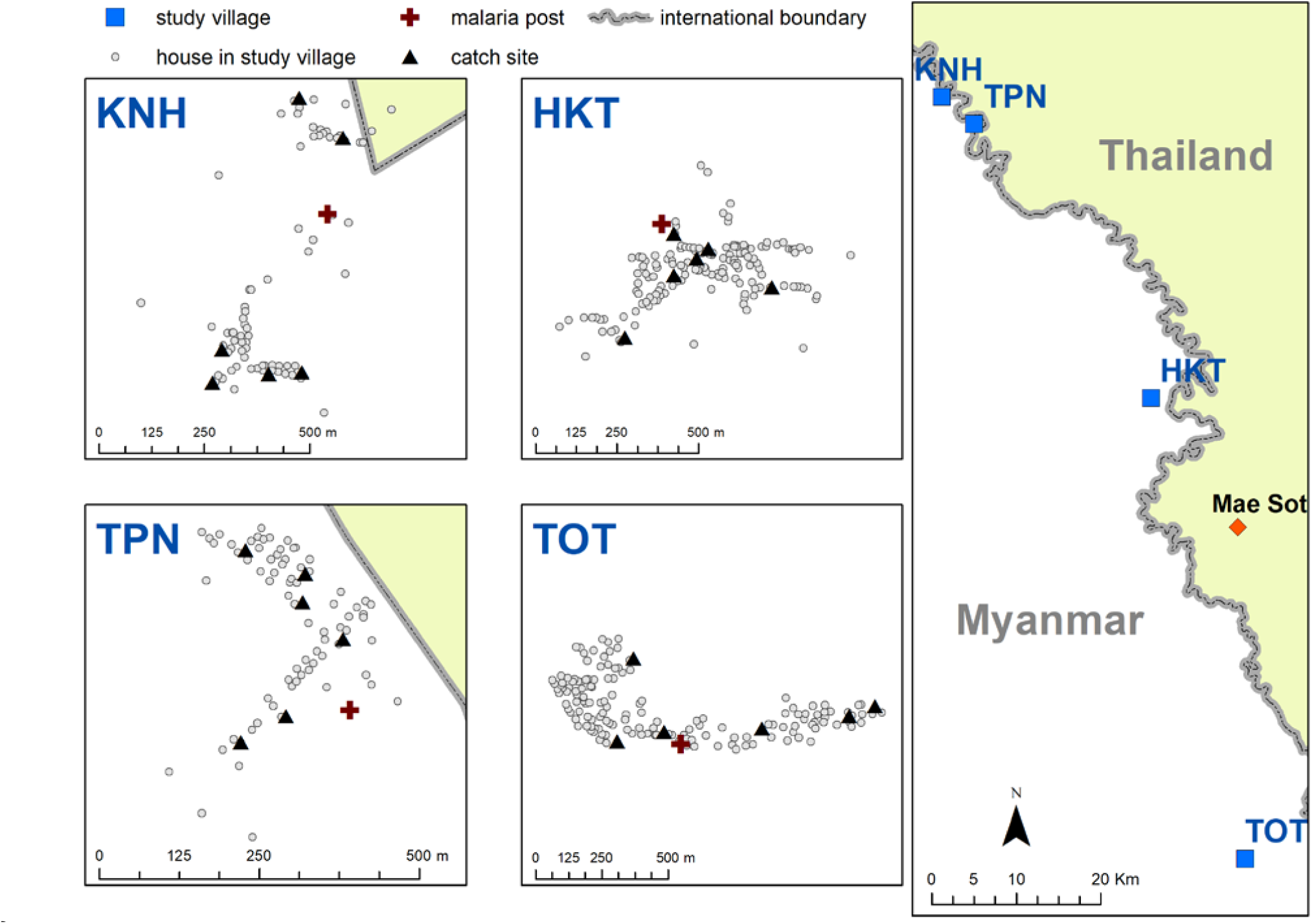
Map indicating the locations of the study villages along the Myanmar-Thailand border; and the distribution of houses, mosquito catch sites and malaria posts within study sites.

A full population census was completed in each of the four study villages at baseline May – June 2013. Everyone enumerated in the census was given a unique identification code. Geographic coordinates were collected for all houses in the four study villages and a unique identification code was assigned to each house. All individuals were then linked to their respective houses.

Blood surveys were conducted at baseline in each village, aiming to screen all individuals above an age of 9 months. Venous blood (3mL) was drawn from each participant, transported to a central laboratory and analyzed using a highly sensitive quantitative PCR (uPCR) assay with a limit of detection of 22 parasites per mL. (Imwong et al., 2014). Subclinical infections detected through these blood screenings are hereafter referred to as uPCR-detected “infections” of either *P. falciparum* or *P. vivax*. Most of these infections (86%) were asymptomatic (J. Landier et al., 2017).

A community-based malaria clinic (malaria post or MP) was established in each village at the beginning of the project. Villagers were trained to diagnose malaria using rapid diagnostic tests (RDTs) and to treat RDT positive infections with dose-based on weight and age. The ID code of each participant who self-presented at the MP was recorded, along with RDT results and these cases are hereafter referred to as clinical malaria episodes of either *P. falciparum* or *P. vivax*. Malaria episodes were treated with dihydroartemisinin-piperaquine (DHA+P) for *P. falciparum* and chloroquine for *P. vivax*. Radical cure for *P. vivax* was not provided because the absence of G6PD tests required to prevent hemolysis in G6PDd individuals (Bancone et al., 2014; Chu et al., 2017).

MDAs were initially conducted in four villages selected based on *P. falciparum* prevalence surveys (using uPCR) in the region. MDAs were conducted in two villages at the beginning of the study and extended to the two control villages beginning in M9. Restricted randomization was used to decide which village received early or deferred MDA. MDA consisted of 3 days of DHA+P, with a single low dose of primaquine on the third day, repeated over three months (M0, M1, M2 for the first group and M9, M10, M11 for the control group). Follow-up blood surveys were conducted in each village every third month after M0 until M18. A final full blood survey was completed in each village at M24.

Mosquitoes were collected monthly using human landing catches to estimate the human biting rate (HBR). Mosquito catching teams were based at 5 sites (both indoors and outdoors) within each of the four study villages (total of 20 catch sites) for 5 consecutive nights during the study period M0 through M20. Mosquitoes were caught using glass tubes and later identified morphologically (Ya-umphan et al., 2016).

The locations of MPs, catch sites and village houses are indicated in Figure 1.

## ANALYSIS

### Variables

All individuals recorded in the census with a house address in the four study villages were included in this analysis.

The data were aggregated into one month time steps and individuals were coded with a “1” for any month in which they presented at the village MP and were diagnosed with a *P. falciparum* or *P. vivax* infection. Likewise, individuals who did not have a clinical episode within a given month were coded with a “0” for that respective month. Individuals who were ever diagnosed with a clinical episode or uPCR detected infection (either *P. falciparum* or *P. vivax*) were likewise coded as a “1” for analyses of having ever been detected by uPCR for an infection or having ever had a clinical episode during the study period.

Individual level predictor variables included age group, gender, infection status (not infected = 0; infected =1) and adherence to MDA (number of doses taken). Household level predictor variables included a binary variable for whether or not another household member had a clinical episode and whether or not another household member had a uPCR-detected infection.

The human biting rate (HBR) for primary vectors (*Anopheles minimus s.l.*, *An. maculatus s.l.*, and *An. dirus s.l.*) was calculated for each month and for each catch site. HBR values were then attributed to individuals based on their house location, assigning the HBR from the catch site that was geographically closest to each house.

Neighborhood MDA adherence was calculated as the proportion of people who took no rounds of MDA within 100 meters radius of each house in the study population. This proportion was calculated for each house in the study villages and non-adherence proportions were then attributed to individuals based on the house to which they were attributed.

Both HBR and neighborhood MDA variables were operationalized as tertiles (lowest third, middle third, highest third, indicated in **Supplementary Table 1**).

### Exploratory spatial and temporal analyses

All predictor variables were explored in bivariate analyses. Unadjusted odds ratios were calculated for binary predictors and Wilcox rank sum tests were calculated for continuous variables. Cumulative hazards curves were used to analyze temporal patterns in infections.

uPCR-detected infections (from surveys) and clinical episodes (from the MPs) were mapped at the house level across time. Maps were created for each village and each survey time point (M0 – M24), with clinical episodes aggregated to align with surveys (i.e. M1, M2 and M3 aggregated into M3).

The standard distance deviation (SDD), a two-dimensional version of a normal standard deviation, was used to visually analyze the distribution of clinical *P. falciparum* episodes for each time point in the one village with sufficient *P. falciparum* infections (TOT). SDD are calculated by finding the mean center of all points, weighted by the number of clinical episodes. The SDD is the average distance from mean center (for both the x and the y plane) for all weighted points and is represented by a circular map layer centered on the weighted mean with the standard distance as radius. One SDD was calculated, corresponding to approximately 68% of all points falling inside of the resulting circle.

Scan statistics were used to test for clustering of uPCR-detected infections (*P. falciparum* and *P. vivax*) across survey months; clinical episodes (*P. falciparum* and *P. vivax*) across all months of the study period and MDA non-participation (operationalized as the number of individuals who took none of the three rounds of MDA). The scan statistics used a moving window (a circle) that centered on each point in the village, testing for the relative risk of cases given a population size within the circle in comparison to the risk outside of the circle. The circle increased in size until it included half of the population and then moved to the next geographic reference point. For Plasmodium infections and malaria episodes the space-time discrete Poisson model was used whereas for MDA participation a purely spatial Poisson model was used (Kulldorff, 1997) (as MDAs were completed within a 3 month time period).

### Logistic regression

Mixed effects logistic regressions were used to calculate model adjusted odds ratios (AOR) and confidence intervals for individual, household and neighborhood level risk factors (variables above) for clinical episodes (*P. falciparum* or *P. vivax*) after MDA. These regressions included a random intercept to account for repeated measures within individuals across the study period.

Most *P. falciparum* infections post-MDA occurred in a single village (TOT), therefore regressions for *P. falciparum* infections post-MDA were based on this village alone whereas regressions for *P. vivax* included all 4 study villages. A final set of regressions looked at the odds of *P. falciparum* or *P. vivax* infection being detected by uPCR after MDA.

### Software

Exploratory statistics and regressions were calculated using R (version 3.4.3; https://cran.r-project.org/) and the “epiR”, “lme4”, and “survival” packages. All maps were created using ArcGIS 10.5 (https://www.arcgis.com/). Exploratory spatial data analysis was conducted using ArcGIS 10.5 and SatScan v9.5 (https://www.satscan.org/). The neighborhood participation variable was created using ArcGIS and the Python programming language (version 3.5.2; https://www.python.org/).

### Ethics approval

This project was approved by the Oxford Tropical Research Ethics Committee (OxTREC: 1015-13; April 29, 2013) and the Tak Province Community Ethics Advisory Board (T-CAB).

## RESULTS

3229 villagers (1689 male) were included in this study. During the study period 80 study participants were diagnosed with clinical *P. falciparum* and 216 with clinical *P. vivax*. 201 and 611 participants were found to have *P. falciparum* or *P. vivax* infections respectively by uPCR and 325 uPCR positive participants had Plasmodium infections not identifiable at the species level. Infections that were not identifiable at the species level were not included in these analyses. Total numbers of clinical episodes and uPCR-detected infections were higher than the total number of infected individuals because some participants had multiple infections.

The majority of clinical *P. falciparum* infections occurred in one of the study villages (TOT). 66 out of the 80 participants who had a clinical *P.falciparum* case were from TOT village (3 from HKT, 7 from TPN and 4 from KNH).

39 (49%) of the participants who had a clinical *P. falciparum* episode lived in a house with someone who had a clinical *P. falciparum* episode (unadjusted odds ratio (UOR): 8.1; CI: 5.1 – 12.7). 35 (44%) of the participants who had a clinical *P. falciparum* episode lived in a house with someone who had a uPCR-detected *P. falciparum* infection during the study period (UOR: 1.6; CI: 1.0 – 2.4).

11 of the 80 participants (14%) who had a clinical *P. falciparum* infection during the study period had repeated clinical episodes. *P. falciparum* and *P. vivax* infections were more prevalent in males than females (UOR: 2.0; CI: 1.5 – 2.8 for *P. falciparum* and UOR: 1.7; CI: 1.4 – 2.0 for *P. vivax*).

### Spatiotemporal patterns in uPCR-detected infections, clinical episodes and MDA adherence

*P. falciparum* and *P. vivax* infections were detected in all villages at baseline (**Figures 2 and 3**). These infections were significantly reduced following MDA in all villages. The prevalence of *P. falciparum* infections had reduced in two villages (villages TPN and HKT) prior to MDA. *P. vivax* infections returned in subsequent months in most villages (as expected given that radical cure was not provided), with the exception of village TPN.

**Figure 2:**
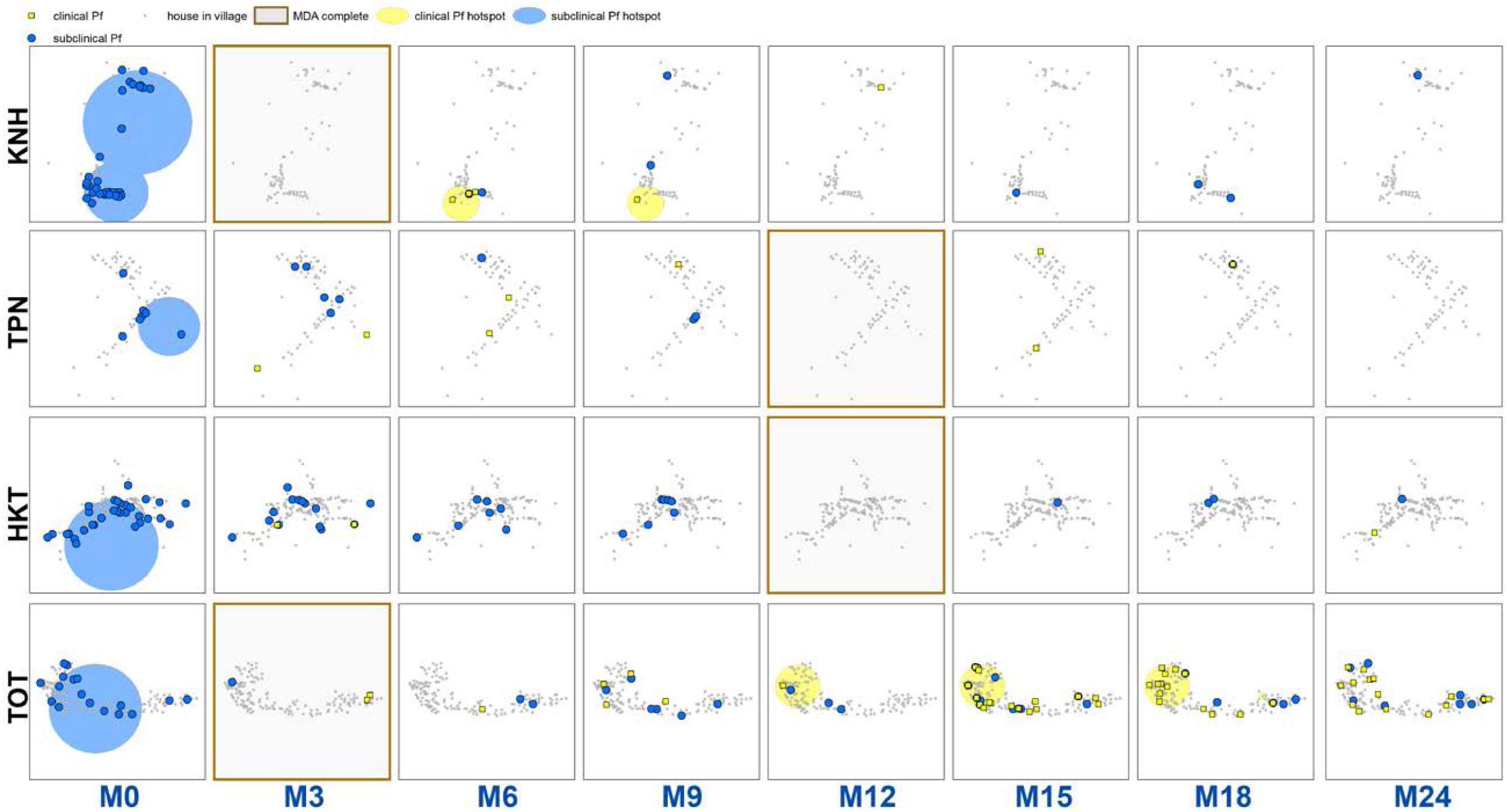
Clinical *P. falciparum* episodes (yellow square points) and uPCR-detected *P. falciparum* infections (blue dots) at house level over time for each of the four study villages. Statistically significant clusters (detected using SaTScan) are indicated for both clinical episodes (underlying yellow circles) and uPCR-detected infections (underlying blue circles).

**Figure 3:**
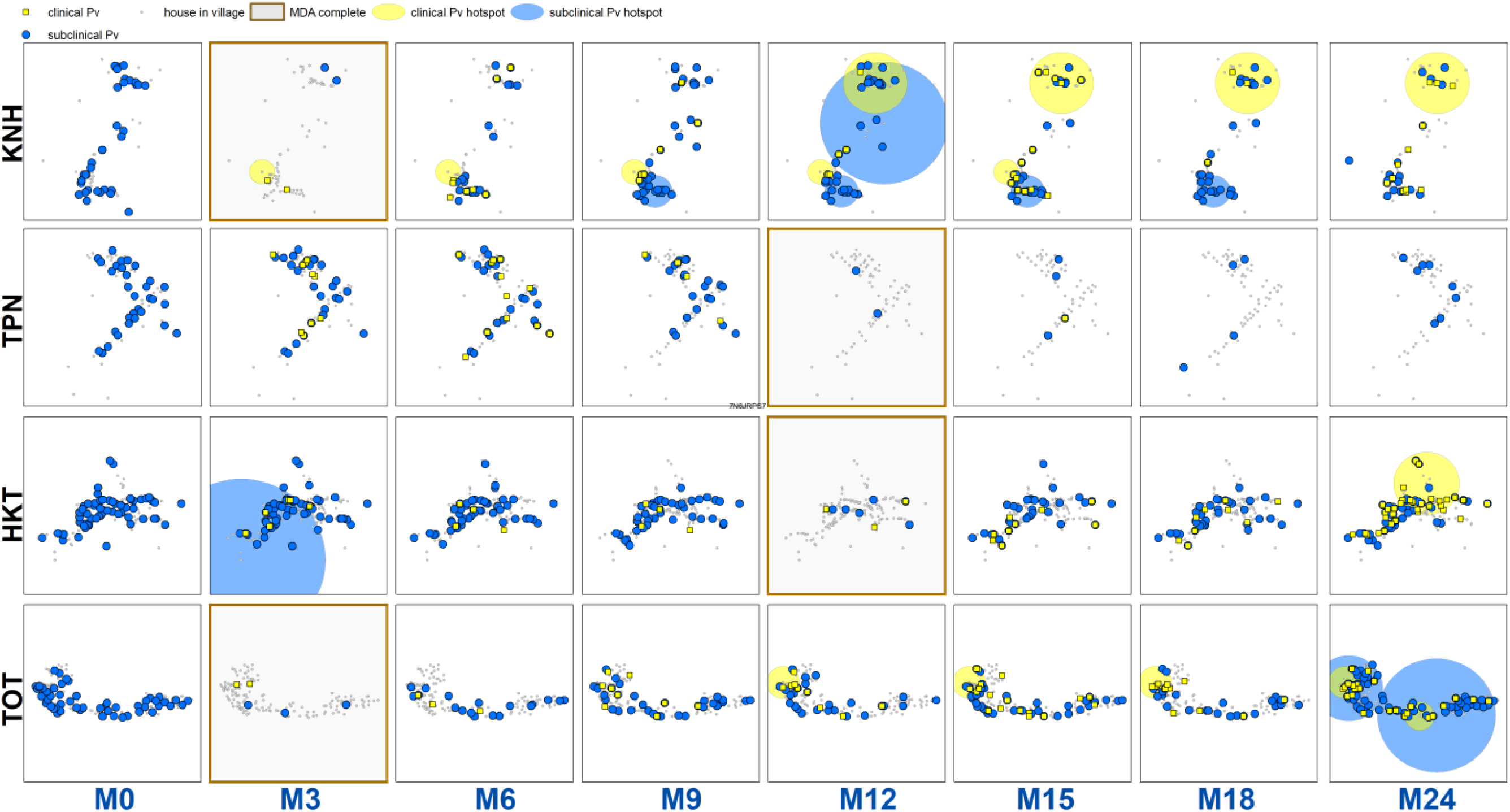
Clinical *P. vivax* episodes (yellow square points) and uPCR-detected *P. vivax* infections (blue dots) at house level over time for each of the four study villages. Statistically significant clusters (detected using SaTScan) are indicated for both clinical episodes (underlying yellow circles) and PCR-detected infections (underlying blue circles).

There were statistically significant clusters of uPCR-detected *P. falciparum* infections in each village at baseline but subsequently no significant clusters were detected (**Figure 2**). Clusters of clinical *P. falciparum* episodes occurred in two villages (KNH and TOT). The cluster in KNH occurred from M5 through M7 but included only four episodes. There were two separate clusters in village TOT. A cluster in the western portion of the village began in M12 and lasted until M18 (with a total of 35 episodes). A single-house cluster occurred in the eastern portion of the village (M15 through M18) with 5 episodes among 4 house members (2 in a 10 yo male, 1 in a 48 yo male, 1 in a 16 yo male, and 1 in a 48 year old female).

*P. vivax* infections were widespread throughout the study villages at baseline but no spatial clustering pattern was detected at M0 (**Figure 3**). Village HKT had a cluster at M3; village KNH had one persistent cluster from M9 through M18; and village TOT had two clusters at M24 (Figure 2). Clusters of clinical *P. vivax* episodes also lingered in village KNH (M3 through M14 (17 episodes) and then M12 through M24 (6 episodes)) and in village TOT (M12 through M21 (31 episodes) and M19 through M23 (18 episodes)). There was a cluster of clinical *P. vivax* episodes in village HKT during M23 and M24 (including 23 episodes).

There were significant clusters of non-participation in the MDAs in three of the study villages (TPN, HKT and TOT (**Supplementary Figure 1**)). The non-participation cluster in TOT made up a large portion of the western part of the village and included 115 individuals not participating in the MDA. The non-participation clusters in HKT and TPN included 206 and 15 individuals respectively.

Sporadic clinical *P. falciparum* episodes occurred in village TOT from M6 to M9, followed by a small outbreak beginning in M13 (**Figure 2**). The first clinical *P. falciparum* episodes during this outbreak occurred among villagers who lived in the cluster of non-MDA participation (**Figure 4**). By M24 the clinical episodes had spread through much of the village (**Figure 4**).

**Figure 4:**
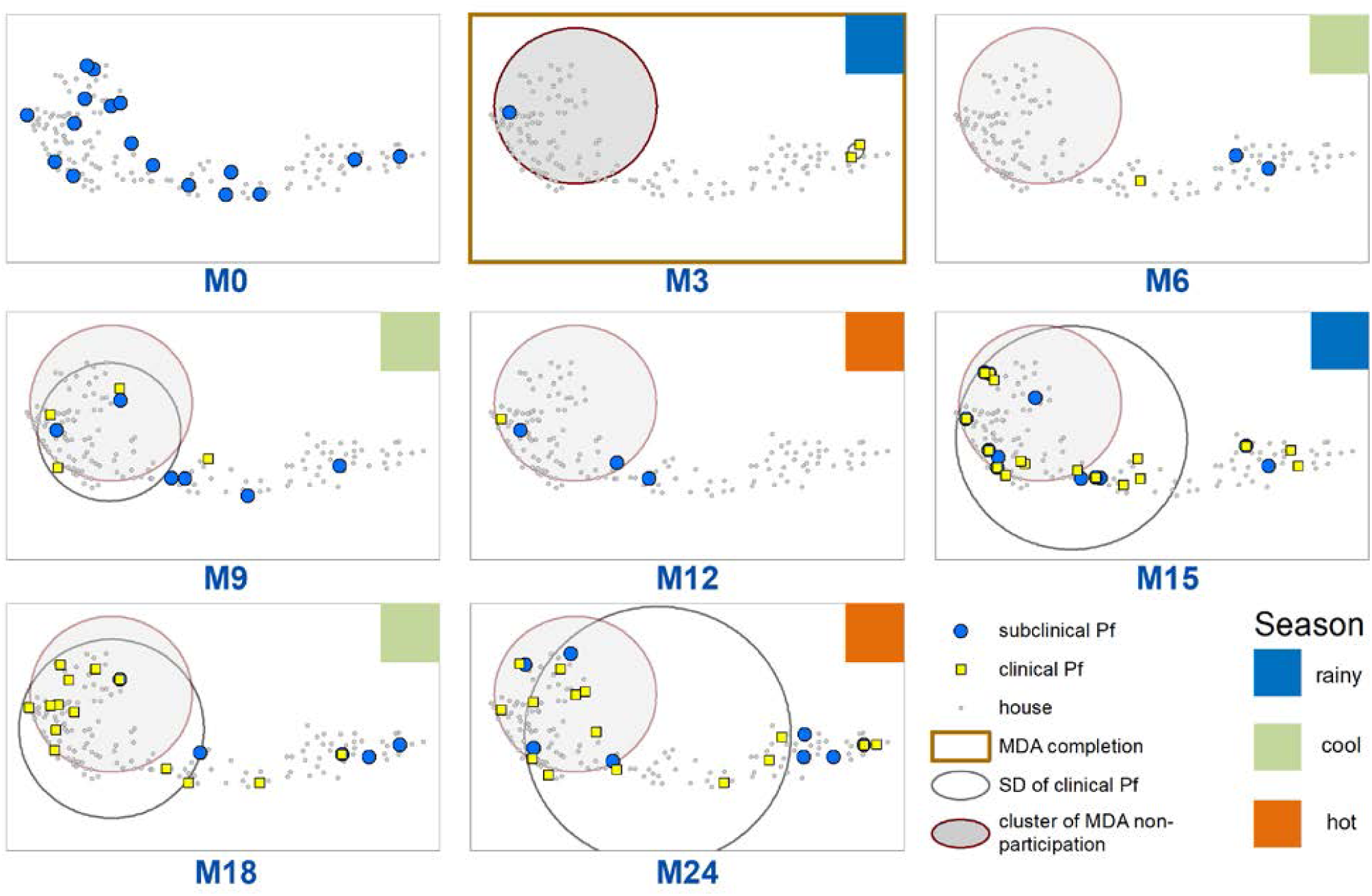
Spatiotemporal distribution of clinical *P. falciparum* episodes (yellow square points), uPCR detected *P. falciparum* infections (blue dots), and a cluster of non-participation in MDA (grey circle/ochre border, detected using SatScan). Season is indicated by colored squares in the top right corner of each map. A measure of the spread of clinical *P. falciparum* cases is given by the standard distance deviation (“SD”), indicated by the hollow circle with dark grey outline. One standard deviation is shown, indicating that roughly 68% of all cases lie inside of the circle. Approximately 6 months post-MDA (M9), clinical infections began in the westernmost portion of the village. The distribution of these cases through month 15 corresponds with the cluster of non-MDA participation (indicated by the grey circle in M3). By M24 the cases extended throughout much of the village.

Cumulative hazards plots of clinical *P. falciparum* episodes in village TOT illustrate the temporal patterns in infections according to neighborhood MDA adherence and household clustering (**Figure 5A**). The proportion of individuals who had acquired a clinical *P. falciparum* episode began consistently increasing in M12 for those living in either low or mid MDA adherence neighborhoods. *P. falciparum* episodes among high MDA adherence neighborhoods began increasing approximately one month after the increase in low adherence neighborhoods but never reached the level experienced in either mid or low MDA adherence neighborhoods. 4.4% of all individuals in high MDA adherence neighborhoods had at least one clinical *P. falciparum* episode by the end of the study period, in comparison to 7.6% in mid and 9.6% in low MDA adherence neighborhoods (log-rank test p-value = 0.0485; **Figure 5A**).

**Figure 5:**
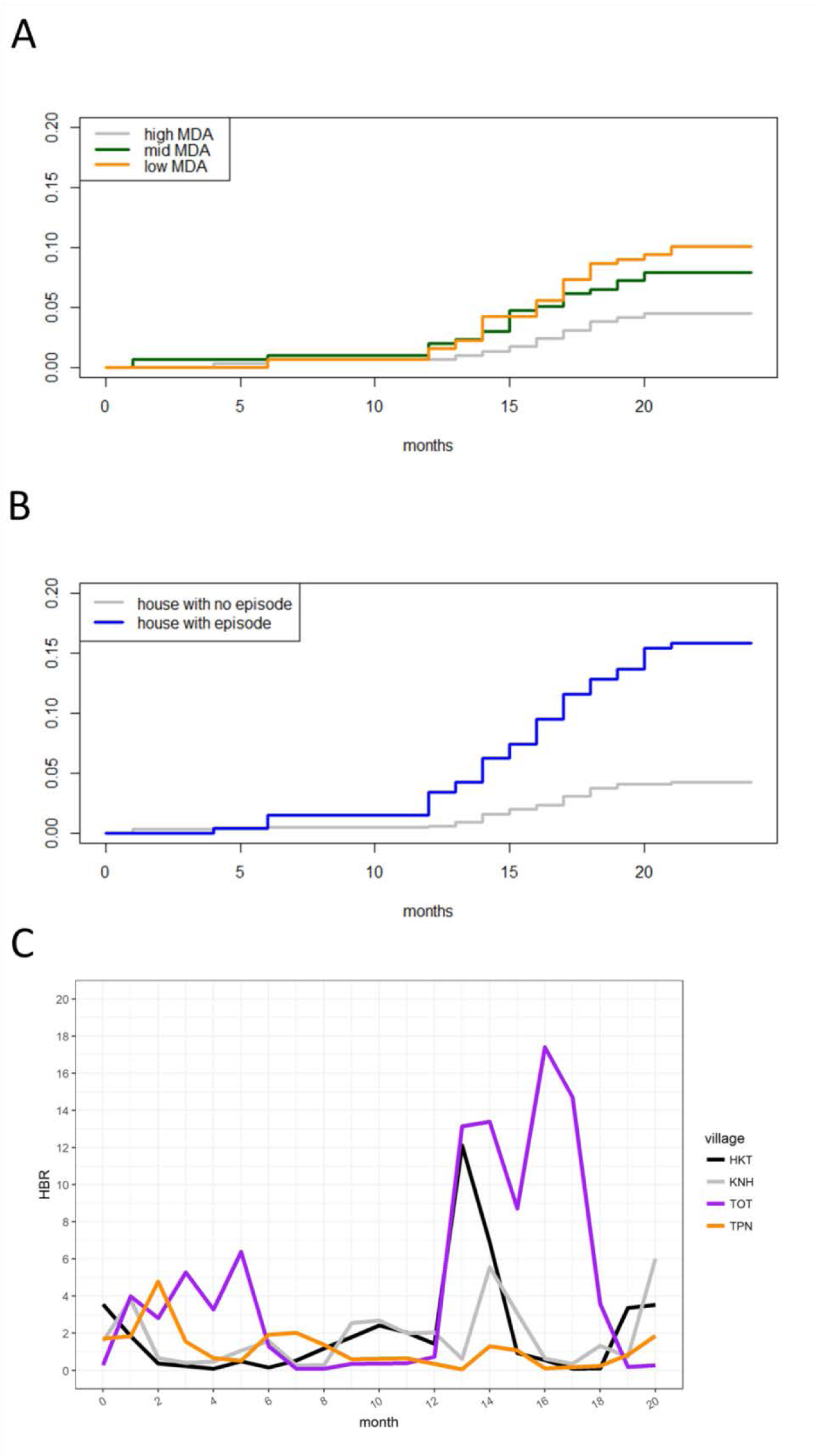
Cumulative hazard for clinical *P. falciparum* episodes in village TOT by A.) neighborhood MDA adherence, B.) household clustering of clinical *P. falciparum* episodes. C.) Indicates the human biting rate (HBR) for primary vectors by study month.

Clinical *P. falciparum* episodes also clustered within houses. Beginning in M6, the proportion of individuals who acquired *P. falciparum* episodes was higher among those who lived in a house with someone else who had a clinical *P. falciparum episode*. The overall proportion of people who acquired a clinical *P. falciparum* episode was consistently higher throughout the study period among those who lived with someone else who had a clinical *P. falciparum* episode (log-rank test p-value < 0.0001; **Figure 5B**).

The increase in clinical *P. falciparum* episodes in M13 coincided with an increase in HBR in village TOT (**Figure 5C**).

### Longitudinal analysis of clinical *P. falciparum* and *P. vivax* episodes

Clinical *P. falciparum* episodes after MDA in village TOT were most likely to occur among 5 to 14 year olds (AOR: 3.0; CI: 1.2 – 7.9) and participants who lived in a house with someone else who had a clinical *P. falciparum* episode during the same month (AOR: 4.2; CI: 2.1 – 8.6) (**Table 3**). Living in a neighborhood with a high proportion of adherence to MDA had a protective effect, being associated with a 90% decrease in the odds of having a clinical episode (AOR: 0.1; CI: 0.01 – 0.51) compared to people who lived in low adherence neighborhoods (**Table 1**). These effects remained after controlling for time.

**Table 1:** Mixed effects logistic regression for odds of having a clinical *P. falciparum* episode (left panel, village TOT only) or clinical *P. vivax* episode (right panel, all four study villages). The models included a random intercept for individual participants, with repeat observations occurring within individuals over the study period.

|  | *P. falciparum* |  |  | *P. vivax* |  |
| --- | --- | --- | --- | --- | --- |
| covariate | AOR | p-value |  | AOR | p-value |
| 0 to 4 | comparison |  |  | comparison |  |
| 5 to 14 | 3.0 (1.17 - 7.90) | 0.0222 |  | 1.0 (0.40 - 2.32) | 0.9433 |
| 15 plus | 1.9 (0.73 - 4.72) | 0.1979 |  | 0.4 (0.17 - 0.99) | 0.0484 |
| female | comparison |  |  | comparison |  |
| male | 1.2 (0.65 - 2.08) | 0.6201 |  | 0.8 (0.41 - 1.53) | 0.4790 |
| no house member with clinical episode | comparison |  |  | comparison |  |
| house member with clinical episode | 4.2 (2.05 - 8.60) | < 0.0001 |  | 5.8 (3.39 - 9.93) | < 0.0001 |
| doses of MDA complete | 0.9 (0.82 - 1.07) | 0.3320 |  | 1.1 (0.90 - 1.28) | 0.4183 |
| low MDA | comparison |  |  | comparison |  |
| mid MDA | 0.4 (0.12 - 1.06) | 0.0630 |  | 0.8 (0.16 - 3.61) | 0.7237 |
| high MDA | 0.1 (0.01 - 0.57) | 0.0100 |  | 0.5 (0.05 - 5.85) | 0.6028 |
| low HBR | comparison |  |  | comparison |  |
| mid HBR | 1.0 (0.42 - 2.33) | 0.9747 |  | 1.6 (1.07 - 2.40) | 0.0210 |
| high HBR | 2.3 (1.17 - 4.46) | 0.0153 |  | 1.6 (1.10 - 2.40) | 0.0147 |
| KNH |  |  |  | comparison |  |
| TPN |  |  |  | 0.6 (0.19 - 1.80) | 0.3450 |
| HKT |  |  |  | 0.3 (0.05 - 1.52) | 0.1377 |
| TOT |  |  |  | 0.7 (0.14 - 4.03) | 0.7275 |

Study month was included as a control (a linear specification was used, but polynomial specifications were also tested). An interaction between individual level MDA adherence and neighborhood level adherence was also included as a control.
The human biting rate (HBR) was operationalized as a time-varying covariate with a one month lag. The covariate for having a house member with a clinical episode was also operationalized as a time-varying covariate (coded as “1” for any month in which another house member had a clinical *P. falciparum* episode and a “0” if no other house members had a clinical episode during that month.)

Living in part of the village with a high HBR was also associated with increased odds of *P. falciparum* infection. Individuals who lived in high HBR portions of the village had over two times the odds of acquiring a clinical episode (AOR: 2.3; CI: 1.2 – 4.5) when compared to those who lived in parts of the village with a low HBR (**Table 3**).

Clinical *P. vivax* episodes also exhibited household clustering. Individuals who lived in a house with someone who had a clinical *P. vivax* episode during the same month had over 5 times the odds of having a clinical *P. vivax* episode when compared to those who did not live in a house with someone who had a clinical *P. vivax* episode (AOR: 5.8; CI: 3.4 – 9.9) (**Table 3**). Living in part of a village with mid or high HBR was also associated with an increased odds of *P. vivax* infection (mid HBR – AOR: 1.6; CI: 1.1 – 2.4; high HBR – AOR: 1.6; CI: 1.1 – 2.4).

### Logistic regression for odds of having a uPCR-diagnosed infection after MDA

The strongest predictor of having a uPCR detected *P. falciparum* infection after MDA was also having a clinical *P. falciparum* episode after MDA (AOR: 4.6; CI: 1.8 – 11.1) (**Table 2**). uPCR detected *P. vivax* infections occurred mostly in older children (AOR: 2.6; CI: 1.7 – 4.1), adults (AOR: 2.3; CI: 1.6 – 3.6) and males (AOR: 1.8; CI: 1.4 – 2.2) (**Table 2**). Individuals who had a clinical *P. vivax* episode after MDA had 2.7 times the odds (CI: 1.9 – 3.9) and individuals who lived in a house with someone else with a uPCR detected *P. vivax* infection had 1.5 times the odds (CI: 1.2 – 2.0) of also having a uPCR detected infection after MDA (**Table 2**).

**Table 2:** Logistic regression for the odds of having a uPCR detected *P. falciparum* (left panel) or *P. vivax* infection (right panel) after MDA. Individuals in the data were coded as having an infection of either species if they were ever determined by uPCR to have an infection through blood screenings in full village blood surveys after MDA. Almost all *P. falciparum* episodes occurred in a single village (TOT) and the analysis for *P. falciparum* was only conducted on data from that village.

|  | *P. falciparum* |  |  | *P. vivax* |  |
| --- | --- | --- | --- | --- | --- |
| covariate | AOR | p-value |  | AOR | p-value |
| 0 to 4 | comparison |  |  | comparison |  |
| 5 to 14 | 4.8 (0.85 - 89.92) | 0.1451 |  | 2.6 (1.74 - 4.08) | < 0.0001 |
| 15 plus | 4.9 (0.96 - 90.57) | 0.1273 |  | 2.3 (1.57 - 3.57) | < 0.0001 |
| female | comparison |  |  | comparison |  |
| male | 1.2 (0.55 - 2.57) | 0.6748 |  | 1.8 (1.42 - 2.24) | < 0.0001 |
| no clinical episode | comparison |  |  | comparison |  |
| clinical episode | 4.6 (1.83 - 11.05) | 0.0008 |  | 2.7 (1.85 - 3.94) | < 0.0001 |
| no house member with uPCR infection | comparison |  |  | comparison |  |
| house member with uPCR infection | 1.2 (0.45 - 2.87) | 0.7226 |  | 1.5 (1.17 - 1.98) | 0.0017 |
| no house member with clinical episode | comparison |  |  | comparison |  |
| house member with clinical episode | 1.4 (0.58 - 3.23) | 0.4518 |  | 1.2 (0.96 - 1.58) | 0.0991 |
| doses of MDA complete | 0.9 (0.65 - 1.00) | 0.2470 |  | 0.9 (0.88 - 1.01) | 0.1160 |
| low MDA | comparison |  |  | comparison |  |
| mid MDA | 0.4 (0.09 - 1.55) | 0.2159 |  | 1.1 (0.67 - 1.82) | 0.7083 |
| high MDA | 0.4 (0.08 - 1.65) | 0.2407 |  | 1.5 (0.70 - 3.15) | 0.2983 |
| KNH |  |  |  | comparison |  |
| TPN |  |  |  | 1.1 (0.71 - 1.58) | 0.7866 |
| HKT |  |  |  | 1.3 (0.75 - 2.38) | 0.3284 |
| TOT |  |  |  | 5.2 (2.84 - 9.54) | < 0.0001 |

The number of surveys that an individual participated in was included as a control variable. An interaction control variable for individual level MDA adherence and neighborhood adherence was also included in the models.

## DISCUSSION

The primary goal of targeted MDA in the study villages was to reduce the prevalence of uPCR-detected *P. falciparum*. There was also a significant impact on the incidence of clinical *P. falciparum* episodes, also as a result of MDA. This analysis shows that post-MDA clinical *P. falciparum* infections in one village, TOT exhibited strong spatiotemporal clustering within houses in low MDA-adherence neighborhoods and within portions of the village with high HBR.

There was no protective effect beyond the prophylactic period after taking MDA at the individual level, but there was a group level effect suggesting a level of herd protection. Those who lived in neighborhoods with high participation in MDA had a reduced risk of becoming infected, regardless of their individual participation in MDA. This protective effect was most pronounced in the rainy season following MDA (**Figure 4**) and corresponded to a surge in vector activity (**Figure 5C**). Once the *P. falciparum* outbreak began, there was a lag of approximately 1 month between the onset of clinical episodes in neighborhoods with mid and low MDA adherence and the spread to neighborhoods with high MDA adherence (**Figure 5A**). However, neighborhoods with high MDA adherence never experienced the same levels of infection as those with mid or low MDA adherence (**Figure 5A, Table 1**). To our knowledge, this is the first documentation of a herd effect conferred of MDA for *P. falciparum* malaria.

HBR was also strongly predictive of clinical *P. falciparum* episodes in village TOT. The increase in clinical *P. falciparum* episodes post-MDA occurred both in areas where MDA adherence was poor and where HBR was high. HBR also peaked in one other village (HKT) at the same time as in village TOT (**Figure 5C**), but occurred in the absence of a detectable parasite reservoir (**Figure 2**) and the HBR did not persist at high levels. Evidence suggests that the MP in TOT was not functioning well in the first year of the study (reported in (J. Landier et al., 2017)) and this influenced the outbreak in TOT.

The combination of a persisting parasite reservoir and persistently high HBR (from M13 – M18) in TOT likely explains the drastically different results between the study villages with regard to *P.falciparum* malaria elimination (**Figure 2**). A better functioning MP in TOT would have reduced the size of the outbreak.

Post-MDA clinical *P. vivax* episodes exhibited spatiotemporal clustering within houses and within areas with mid to high levels of HBR. uPCR-detected *P. vivax* infections also clustered within houses. While there was an immediate reduction in blood-stage *P. vivax* following MDA (evident in **Figure 3**) there was no overall effect of MDA on the risk of subsequent clinical episodes or uPCR-detected infections over the entire surveillance period (Chaumeau et al 2018; not yet published).

Clinical *P. vivax* episodes occurred more commonly among younger age groups while uPCR-detected infections occurred more commonly among adults. This pattern was previously described in Thai-Burma border populations over two decades ago (Luxemburger et al., 1996) and is likely the result of some level of acquired immunity to *P. vivax* infections with time. Both adults and children are exposed to similar levels of *P. vivax* transmission, yet in children the infection is more likely to result in clinical symptoms whereas in adults, many of the infections are likely to become or remain asymptomatic.

Clustering of *P. falciparum* infections across houses occurred for limited periods of time only prior to MDA (**Figure 2**). Clustering of *P. vivax* across houses persisted across time before and after MDA (**Figure 3**). Overlapping clusters of clinical *P. vivax* episodes and uPCR-diagnosed *P. vivax* infections were observed in KNH (M9 through M15) and TOT (M24). There were overlapping clusters of clinical P. vivax episodes and clinical *P. falciparum* episodes in village KNH (M5 – M7) and village TOT (M12 – M18). The spatiotemporal clustering patterns of both *P. falciparum* and *P. vivax* suggest that interventions such as reactive case detection would have resulted in the detection of extra cases (of both clinical and uPCR-detected *P. falciparum* and *P. vivax*) when searching within houses and occasionally in neighboring houses, but these would have only been a small proportion of all infections within the villages (Parker et al., 2016) and would not have halted transmission. Conversely, community based EDT and MDA with high participation, targeted at the village scale or larger, appear effective at reducing prevalence, incidence and transmission of *P. falciparum* (Jordi Landier et al., 2018).

Individuals with clinical episodes of both *P. falciparum* and *P. vivax* were frequently, diagnosed (either before or after experiencing an episode) with asymptomatic infections during blood screenings. The study did not genotype the infections. In this low transmission setting at least a proportion of these associated clinical episodes and uPCR detected infections are likely to have been the same parasite strain, suggestive of long-term carriage of both *P. falciparum* and *P. vivax*. This is supported by a large Vietnamese cohort study, showing that 20% of asymptomatic *P. falciparum* carriers and 59% of *P. vivax* carriers carry their parasitaemia for 4 months or longer (Chen et al., 2016; Nguyen et al., 2018). The observation that *P. falciparum* and *P. vivax* parasite densities oscillate during long-term Plasmodium carriage suggests that gametocytaemia might also occur in waves, at some point in time sufficient to transmit to a suitable vector. If such infections remain asymptomatic, they are unlikely to be diagnosed and treated through standard early diagnosis and treatment (EDT) approaches. Active approaches for detecting and treating these infections, such as targeted MDA or mass screen and treat, are necessary.

There are several limitations to this work. Individuals who did not participate in MDA also did not participate in blood screenings immediately after MDA (i.e. M3 in village TOT). uPCR detected infections are therefore likely to be underdiagnosed for these individuals. There is also evidence of a poorly functioning MP in this village, which could have meant undiagnosed (and therefore unrecorded) clinical episodes, especially during the beginning of the study. Finally, some infections are likely to be acquired outside of the village, leading to complex spatial patterns in infections that are mapped at the house level. Within household clustering can be the result of within house transmission, or shared exposure among household members.

## CONCLUSION

These data suggest that poor MDA adherence in the presence of a significant parasite reservoir and sufficient vector activity, can lead to resurgence of malaria in a matter of months after reduced transmission due to MDA (**Figure 5A**). This is especially true in a setting with a poorly functioning diagnosis and treatment center. Conversely, high coverage of MDA in a population can confer benefits long after the prophylactic period has ended – most likely as a result of clearing the parasite reservoir (**Table 1**, **Figure 5A**). Community engagement to maximize community participation, is crucial for MDA success (Kajeechiwa et al., 2017) and villages or portions of villages with poor adherence present persistent challenges to elimination efforts.

## ACKNOWLEDGEMENTS

We would like to thank the study communities in Kayin State, Myanmar for their participation, support and acceptance. We would also like to acknowledge the many staff members at Shoklo Malaria Research Unit and the Mahidol-Oxford Tropical Medicine Research Unit who made this project possible. This study is part of the larger “Targeted Chemo-elimination (TCE) of Malaria (TME)” project, which is registered at ClinicalTrials.gov: NCT01872702 (https://clinicaltrials.gov/ct2/show/NCT01872702). Funding for the TME project was obtained from Wellcome Trust (101148/Z/13/Z) to Prof. Nicholas J. White and the Bill and Melinda Gates Foundation (OPP1081420) to Prof. Arjen M. Dondorp. Sai Thein Than Tun is supported by the Wellcome Trust (grant no. 205240/Z/16/Z).

---

**Supplementary Figure 1:**
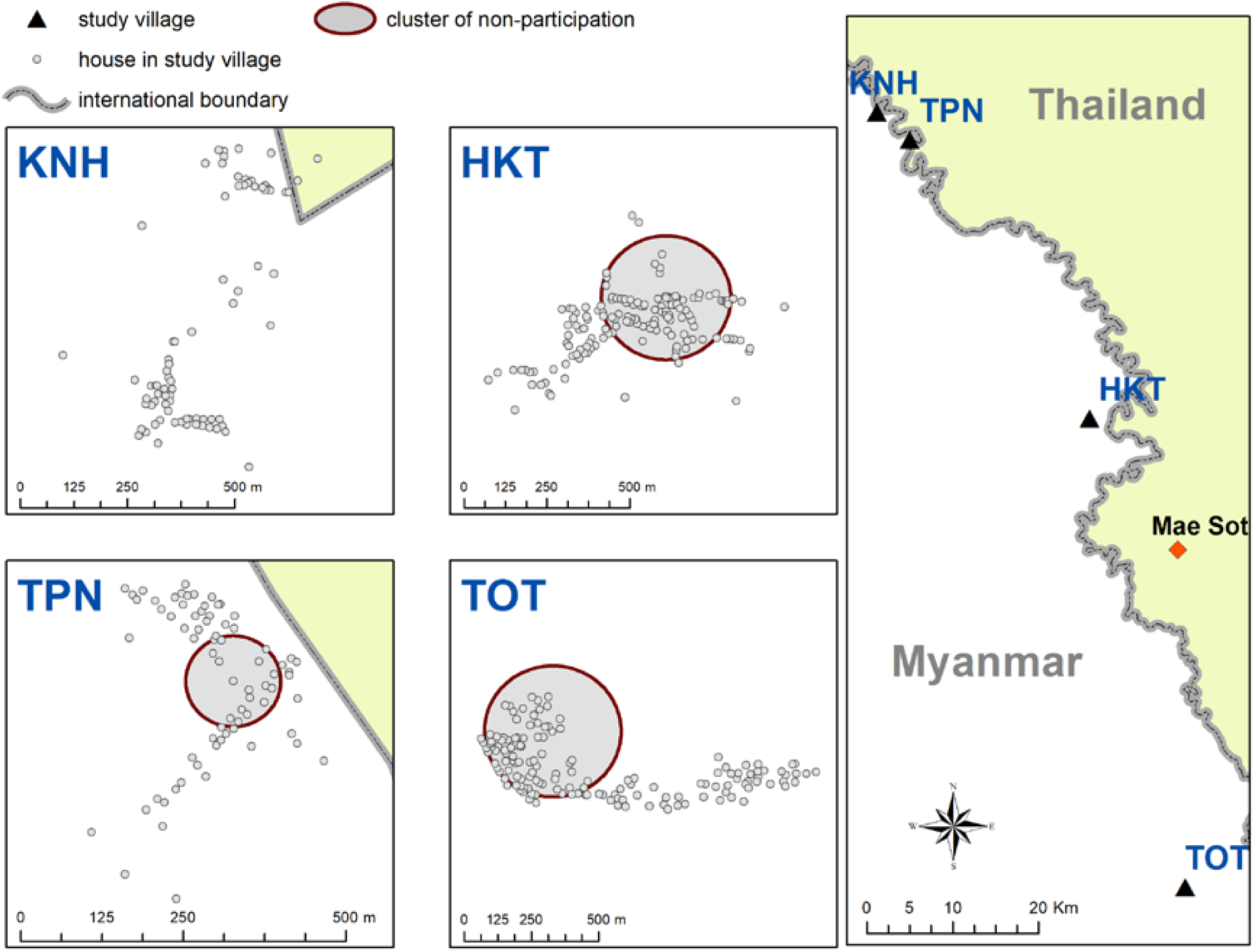
Spatial clusters (detecting using SatScan) of non-participation in MDA

**Supplementary Table 1:** Tertiles (lower, middle and upper 1/3) of MDA non-adherence (% taking no rounds of MDA) and HBR by all villages (used in regression for *P. vivax*) and for TOT alone (used in regression for *P. falciparum*)

|  | All villages | TOT only |
| --- | --- | --- |
| high MDA | $\geq 0.27$ | $\geq 0.294$ |
| mid MDA | $\geq 0.10$ & $< 0.27$ | $\geq 0.20$ & $< 0.294$ |
| low MDA | $< 0.10$ | $< 0.20$ |
| high HBR | $\geq 1.94$ | $\geq 3.70$ |
| mid HBR | $\geq 0.42$ & $< 1.94$ | $\geq 0.36$ & $< 3.70$ |
| low HBR | $< 0.42$ | $< 0.36$ |

